# Buffered sweat microfluidics with AI-enabled translation for clinically actionable blood urea estimation and renal risk stratification

**DOI:** 10.64898/2026.06.17.732831

**Authors:** Jiahao Zhang, Zhengquan Shen, Ming Xu, Yukun Ge, Xi Ren, Genlin Liu, Xiuwei Zhang, Simian Fu, Cheng Yang, Minghui Long, Shu Li, Gaozhi P. Mo, Yi Gong, Nan Li, Peiyu Ma, Zhiyong Peng, Yunlong Zhao

## Abstract

Kidney-function assessment relies on blood urea as a clinically informative metabolic marker; however, its dependence on venipuncture and centralised laboratory testing limits high-frequency monitoring and delays timely clinical intervention. Here, we report an integrated platform combining a wearable buffered microfluidic patch with a physiology-informed, data-driven calibration framework for real-time, non-invasive estimation of blood urea from microlitre-scale sweat volumes (4.79 μL). By precisely regulating the release kinetics of internal buffer salts, the device stabilises the local reaction microenvironment, mitigating variability in sweat pH and flow to ensure reproducible measurement. The resulting signals are processed through an artificial intelligence (AI)-enabled analysis pipeline that integrates sweat urea with patient-specific physiological information to generate clinically interpretable outputs. In multicentre studies, sweat urea shows a strong association with blood urea across diverse cohorts, but with nonlinear and time-lagged relationships that limit direct use. The AI-enabled calibration model compensates for these effects, enabling high-fidelity estimation of blood urea (r = 0.945 versus gold-standard measurements) at clinically relevant concordance levels. The platform further identifies kidney injury with 89.1% accuracy and stratifies disease severity with 83.2% accuracy. Notably, these results demonstrate that the integration of physicochemical stabilisation and AI-enabled data-driven translation establishes sweat as a clinically actionable surrogate for renal monitoring, supporting population-level estimation and highlighting the potential for personalised longitudinal assessment, and enabling a scalable, non-invasive strategy for high-frequency kidney disease management.

## Introduction

Chronic kidney disease (CKD) affects approximately 850 million people worldwide, representing nearly 10% of the global population and rising as one of the leading causes of mortality^1-3^. A formidable challenge in clinical management is the stark contrast between the insidious progression of early-stage CKD and the abrupt deterioration seen in acute kidney injury (AKI). In early CKD, the kidney’s robust compensatory capacity maintains metabolic balance through hyperfiltration in remaining nephrons, rendering the disease largely asymptomatic and “silent” until late-stage failure^4^. Conversely, in AKI or acute exacerbations, a rapid decline in nephron function triggers a catastrophic “waterfall effect,” where the precipitous accumulation of nitrogenous wastes and electrolyte imbalances causes irreversible renal failure within hours^5-7^. This bi-phasic nature of renal disease, characterised by long-term silence followed by rapid collapse, demands high-frequency monitoring of key metabolic markers to enable early intervention and dynamic treatment titration^4-7^.

Renal function assessment currently relies on blood-based biomarkers, most notably blood urea and serum creatinine^8,9^. While central to clinical guidelines, these metrics are tethered to invasive venipuncture and centralised laboratory infrastructure, requiring sophisticated automated analysers and trained personnel. These requirements impose significant barriers: for high-risk populations, they preclude the frequent screening necessary to detect “silent” early-stage CKD; for hospitalised patients, the low sampling frequency and long sample-to-result turnaround times fail to capture the hyper-acute dynamics of AKI^6,9^. These limitations highlight the need for alternative sensing strategies that are non-invasive and suitable for frequent monitoring, while achieving agreement with gold-standard blood measurements at levels typically required for clinical decision-making (typically around 90%, depending on application) ^10^, ideally in simple and deployable formats that enable rapid and actionable feedback outside traditional clinical settings, analogous to the paradigm-shifting glucose test strip ^11^.

Sweat urea, partitioned from blood via the sweat glands, has emerged as a promising non-invasive surrogate for blood urea monitoring (**Fig.1a, upper left**)^12-16^. However, translating sweat measurements into clinically reliable indicators of renal function remains an unresolved challenge^13,16,17^. This limitation arises from both analytical and physiological instabilities inherent to sweat-based sensing. In current clinical practice, urea is most reliably quantified using enzyme-based assays, including urease-coupled colourimetric methods, which depend on tightly controlled reaction conditions. When applied to sweat, however, assay performance becomes highly sensitive to variations in pH and ionic composition, both of which exhibit substantial interindividual variability^12^. Similar environmental susceptibility is observed across alternative sensing modalities. Furthermore, the low secretion volumes (microlitre-scale) and rapid evaporation of sweat frequently lead to sample contamination and measurement errors^17-20^. More crucially, in dynamic clinical states such as dialysis, the sweat– blood urea relationship is not a simple linear mapping but is confounded by complex nonlinear dynamics and temporal lags, necessitating a more sophisticated framework than traditional one-to-one correlation^21,22^.

Here, we report a buffered microfluidic platform coupled with physiology-informed, data-driven mapping that overcomes these barriers to enable precise, non-invasive blood urea estimation and renal risk stratification (**Fig. 1**). We developed a wearable microfluidic patch that requires only 4.79 μL of sample for real-time on-device urea quantification (**Fig. 1a**). To mitigate interindividual variability in sweat composition, we integrated a phosphate-buffered saline (PBS) kinetic release system that stabilises the reaction microenvironment, as supported by numerical simulations of pH distribution and reaction kinetics. To bridge the disparity between sweat and systemic measurements, we further developed a data-driven calibration model that integrates sweat urea with patient-specific physiological and clinical variables, learned from large-scale multicentre datasets (**Fig. 1b**). Implemented within a smartphone-based analysis pipeline, the system enables automated image acquisition, signal extraction and translation into clinically interpretable outputs. Appropriate microfluidic buffering fully utilizes the available colorimetric range and produces a smooth concentration-dependent response (**Fig. 1c**). By capturing physiological correlations, including temporal lag and nonlinear transport effects, the model enables quantitative mapping of sweat signals to high-fidelity blood urea estimates (r = 0.945 versus gold-standard measurements, **Fig. 1d**). Notably, the platform achieves clinically relevant performance in identifying kidney injury (89.1%) and stratifying disease severity (83.2%) and has demonstrated its feasibility for near-patient testing scenarios in real-world clinical workflows. Together, these results establish buffered sweat microfluidics as a clinically actionable surrogate for blood urea and define a general framework for coupling physicochemical stabilisation with data-driven translation of biofluid signals, enabling non-invasive, high-frequency and personalised kidney disease management.

**Fig. 1.**
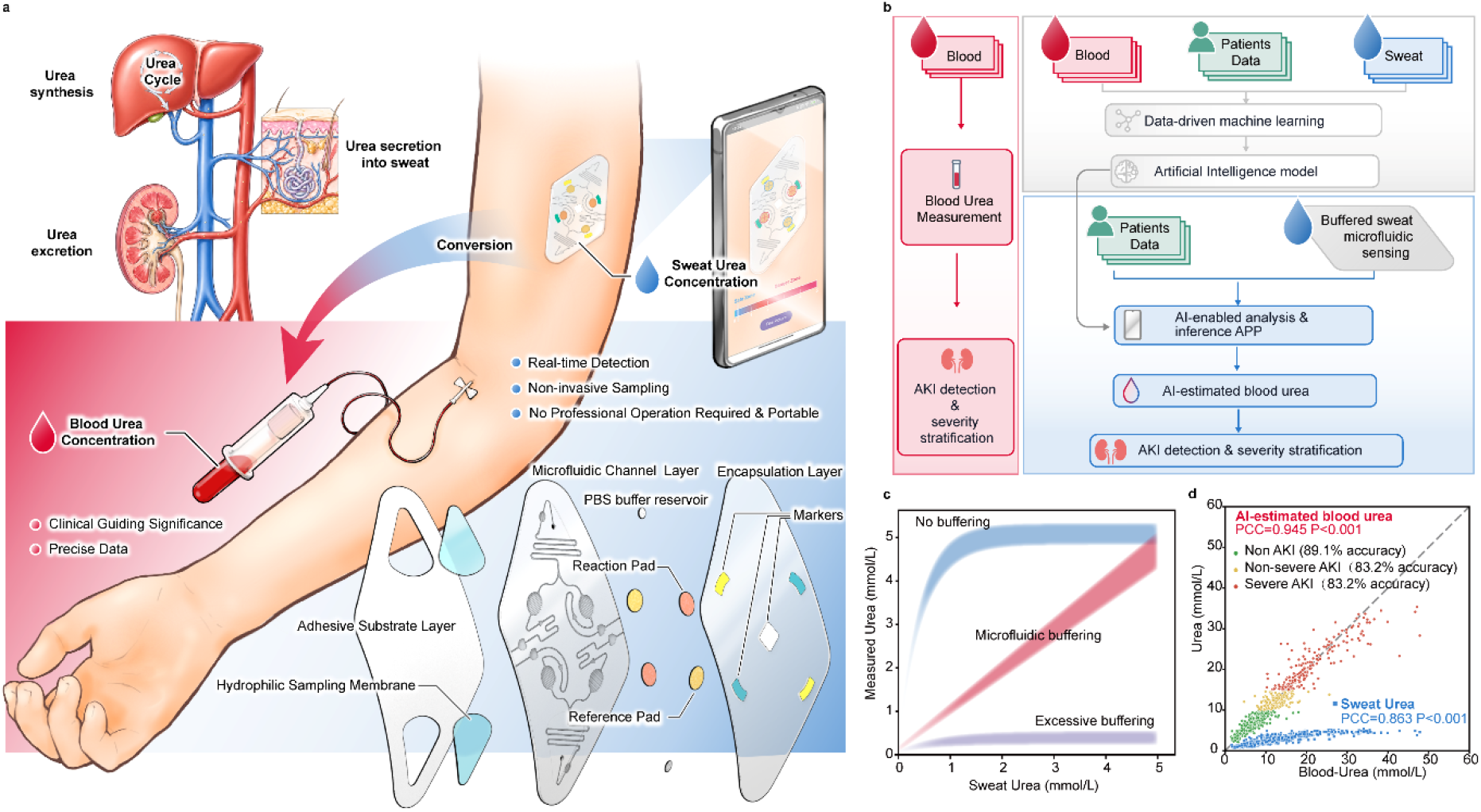
Buffered microfluidic platform with AI-enabled translation for sweat urea sensing and clinical inference. **a)**Schematic of the wearable sensing concept. Urea produced from the urea cycle is partially excreted in sweat (upper left). A microfluidic colourimetric patch enables non-invasive, real-time sweat collection and quantification, with image acquisition and analysis performed through an AI-enabled pipeline to estimate blood urea (bottom). The device architecture (exploded view) comprises an adhesive substrate layer, hydrophilic sampling membrane, microfluidic channel layer and an encapsulation layer, with fiducial colour and registration markers for image normalisation. **b)** End-to-end data pipeline for physiological translation and clinical inference. Sweat urea measurements are integrated with patient-specific physiological and clinical variables for model training; the trained model is subsequently deployed to generate calibrated blood urea estimates and support downstream diagnostic assessment. **c)** Effect of microfluidic buffering on sweat urea sensing. Appropriate buffering expands the dynamic range and enables a smooth concentration-dependent response, whereas insufficient or excessive buffering causes signal saturation or suppression. **d)** Parity plot of estimated versus measured blood urea. Raw sweat urea (blue square) deviates from the identity line (dashed), whereas AI-estimated blood urea from sweat urea concentration (circle) shows improved agreement and more closely follows y = x.

## Results and discussion

### Design and engineering of the buffered microfluidic patch

To achieve reliable sweat urea sensing despite interindividual variability in sweat composition, we first designed a wearable microfluidic patch with integrated on-chip buffering for reproducible measurement. The device architecture (**Fig. 1a**) comprises a medical-grade adhesive substrate layer with integrated sweat reservoir and pilocarpine delivery, a hydrophilic sampling membrane, a microfluidic channel layer and an encapsulation layer, enabling controlled sampling and on-body analysis. The device was fabricated via multilayer PDMS replica moulding, curing and demoulding, followed by reagent loading and plasma bonding (**Fig. 2a**). Because the intrinsic hydrophobicity of the channel material can impede capillary filling, we asked whether selective surface modification could enable reliable fluid transport without compromising device integrity. To this end, the enclosed channels were rendered hydrophilic post-assembly by flowing a silane reagent through the microchannels, reducing the water contact angle from 118° to 48°, and enabling efficient capillary transport while maintaining stable interlayer bonding.

**Fig. 2.**
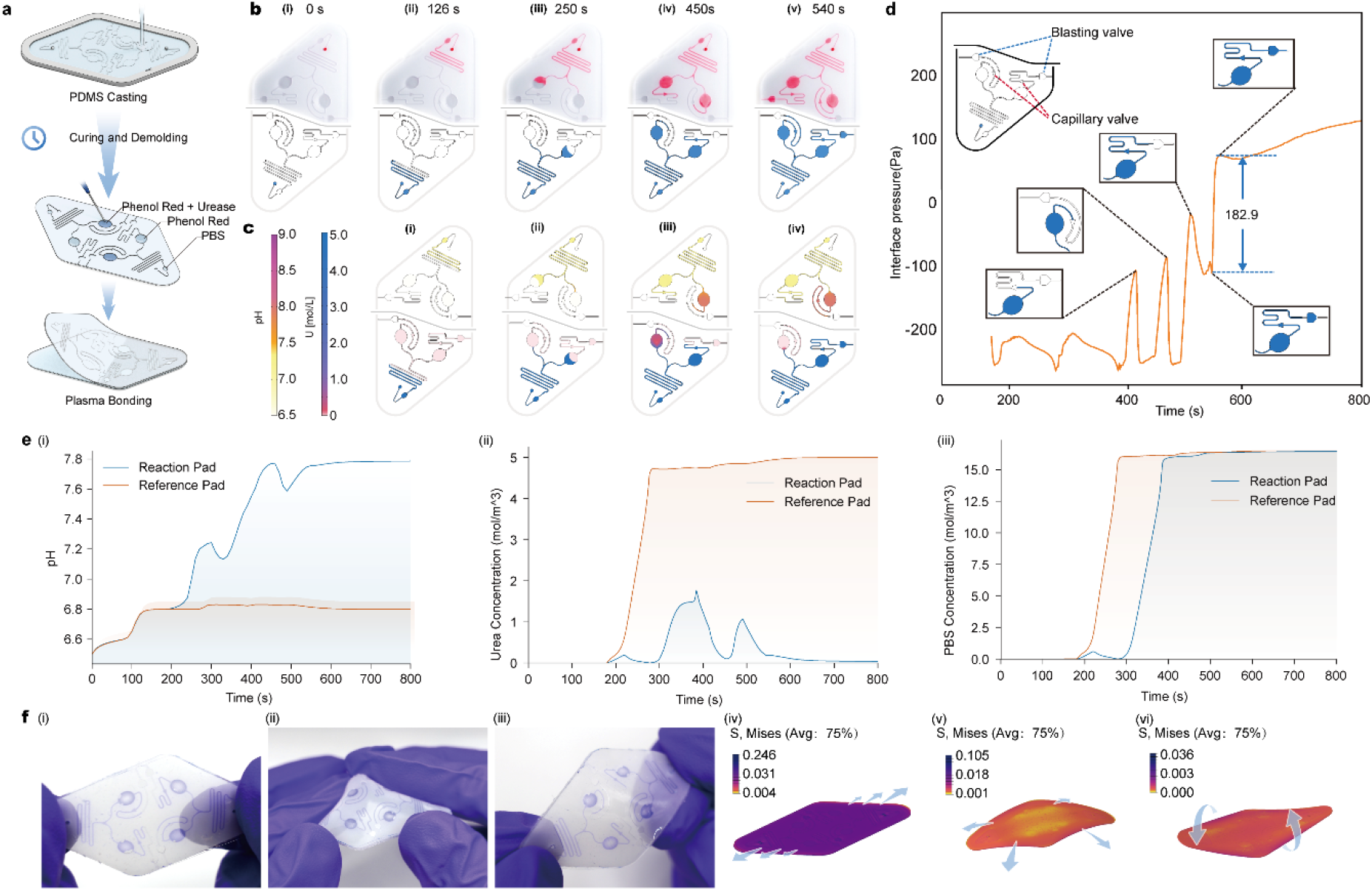
Design and characterisation of the microfluidic system. **a)**Schematic of device fabrication and assembly, including PDMS casting, curing, demoulding, reagent preloading (phenol red with urease in the reaction pad, phenol red in the reference pad and PBS in the buffer pad) and plasma bonding. **b)** Comparison of capillary filling behaviour and fluid–air interface evolution in injection experiments (top) and phase-field simulations (bottom) at 0.45 µL min^−1^, shown at the indicated times. **c)** Multiphysics simulations of coupled transport–reaction during filling at 0.45 µL min^−1^, showing spatial maps of pH (top) and urea concentration (bottom). **d)** Time evolution of the liquid–gas interfacial pressure in the reference and reaction branches. The dashed line indicates the burst-pressure threshold. Once the pressure in the reference branch reaches the threshold, the reference-side valve opens preferentially, releasing excess fluid while retaining the reaction volume in the reaction pad. **e)** Time evolution of pad-averaged quantities in the reaction pad (purple) and reference pad (blue): (i) pH, (ii) urea concentration, and (iii) PBS concentration. **f)** Mechanical characterisation of the device: Representative photographs (right) and finite-element analysis of deformation (left) under (i,iv) 10% uniaxial tension, (ii,v) bending over a 50-mm-radius form, and (iii,vi) 40% torsional shear.

To further improve measurement reliability under physiological variability, we incorporated an upstream PBS buffer reservoir and paired phenol red sensing pads, with urease localised exclusively in the reaction pad to enable differential readout (**Fig. 1a**). Because the device requires only a microlitre-scale volume (∼4.79 µL) to fully fill the microfluidic network, enabling operation even under low sweat production, we next evaluated whether capillary-driven flow could reliably fill the branched channels. Benchtop experiments using a dyed solution infused at 0.45 µL min^−1^ showed reproducible, sequential filling of the microchannels (**Fig. 2b**). These dynamics were reproduced by a two-phase computational fluid dynamics (CFD) model describing liquid–air interface evolution (**Fig. 2c**), with agreement between simulated and measured volumetric filling profiles. The liquid first wetted the buffer pad and serpentine mixer, then split at the Y-junction to fill the reference and reaction pads. The arrow-shaped capillary check valves at the two pad outlets pin the advancing menisci, allowing pressure to build up and ensuring complete wetting of both pads. Subsequently, the menisci break through the arrow-shaped capillaries and continue moving toward the outlet, where they are pinned by burst valves near the outlet. When the interfacial pressure difference reaches approximately 182.9 Pa, the burst valve on the reference pad side opens, releasing excess liquid through the reference branch, while the valve on the reaction side remains locked, thereby confining the reaction volume within the pad and minimising flushing effects (**Fig. 2d**).

We next asked whether on-chip buffering could establish a stable chemical microenvironment for enzymatic sensing. A coupled transport–reaction model incorporating diffusion, acid–base equilibria and urease-catalysed urea hydrolysis predicted that key chemical variables, including local pH, urea concentration and buffer distribution, evolved dynamically but approached steady state within ∼500s, while urea consumption in the reaction pad was completed by ∼600 s (**Fig. 2e**). Consistent trends were observed for carbon- and nitrogen-containing species, which similarly approached steady state within ∼500 s. These results indicate that the buffering system establishes a quasi-stable chemical environment prior to the sensing endpoint, supporting robust colourimetric readout.

We next assessed mechanical robustness under skin-relevant deformation conditions. Finite-element analysis (**Fig. 2f i–iii**) showed that representative modes of wearable deformation, including stretching, bending and twisting, induced a maximum Von Mise stress ∼0.246 MPa within the device, remaining well below the failure limits of the laminated structure (0.51 MPa)^23^. Consistent with these predictions, experimental testing (**Fig. 2f iv–vi**) demonstrated that the device maintained structural integrity under these deformations without leakage or delamination, supporting conformal wear during practical use.

### Physiology-adaptive buffering and integrated colorimetric readout for non-invasive urea quantification

While the microfluidic system ensures reliable fluid transport and mechanical robustness, accurate biochemical sensing requires a controlled reaction microenvironment that can be disrupted by physiological variability in sweat composition. Urea quantification relies on urease-catalysed hydrolysis, which increases local pH and is transduced into a colorimetric signal by phenol red. However, substantial variations in sweat pH across individuals can alter the initial reaction conditions and introduce bias in urea readout. In addition, deviations in the initial pH can shift the system outside the effective operating window of phenol red (pH transition range: 6.8–8.2), resulting in signal saturation and loss of dynamic range, thereby limiting reliable urea quantification.

To mitigate this variability and implement on-chip pH conditioning, we designed a structured PBS buffer pad upstream to regulate buffer release and pre-condition incoming sweat. A reduced transport model was used to relate geometric parameters of the groove structure, including pitch (p) and porosity (ϕ), to an effective buffer delivery rate (k_eq_), thereby mapping device structure to buffer flux (**Fig. 3a**). The optimised groove geometries were implemented in the device, as confirmed by microscopy images showing well-defined and spatially patterned microchannels (**Fig. 3b**), which regulate buffer–fluid interaction and hence buffer delivery into the flowing stream.

**Fig. 3.**
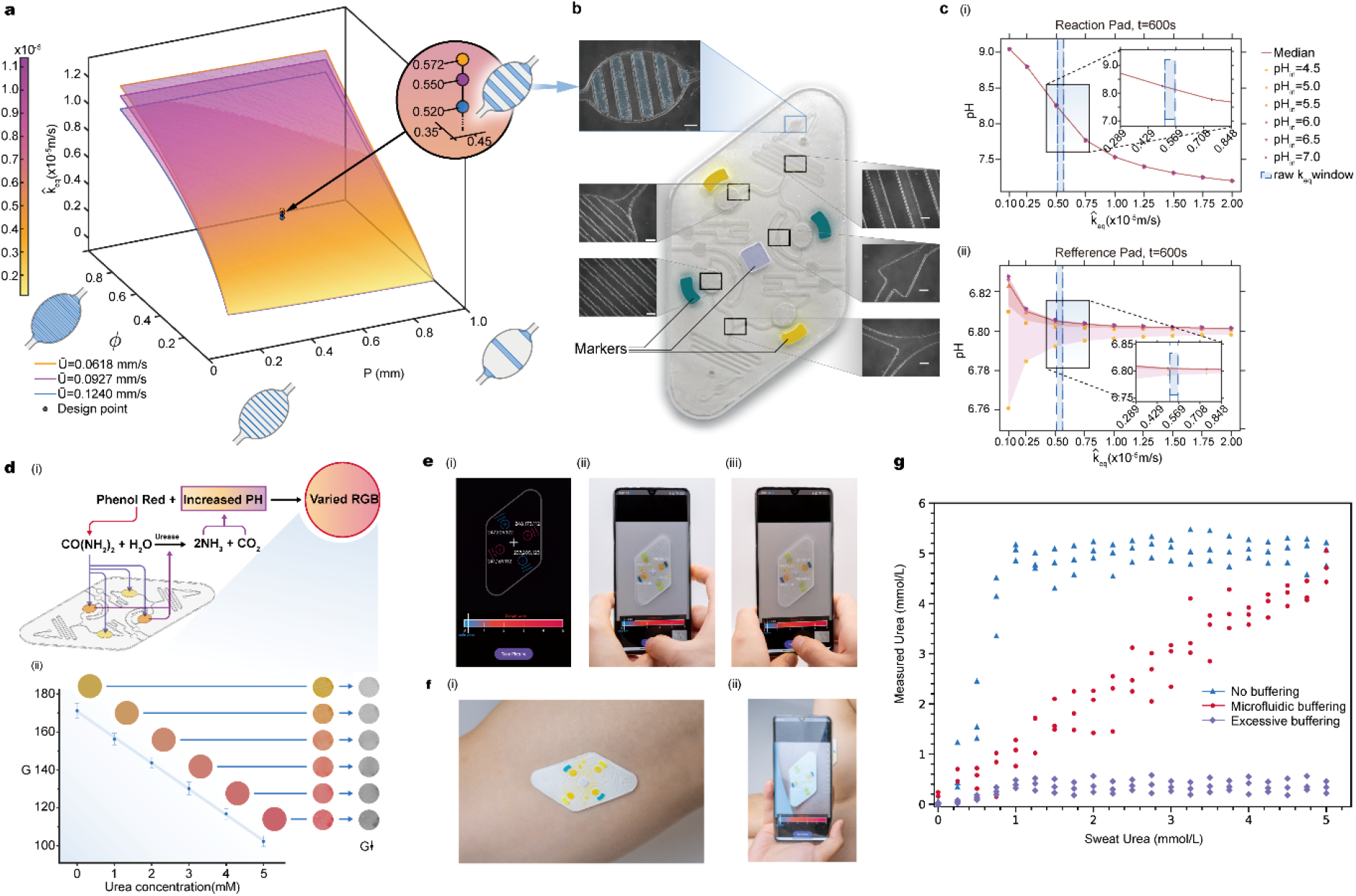
Design and characterisation of the pH-buffering system against physiological difference and colorimetric analysis. **a)**Derived equivalent surface mass-transfer coefficient (*k*_*eq*_) as a function of groove design parameters (pitch *p* and porosity *ϕ*) for the indicated mean flow velocities (*U*). **b)** Optical microscopic photograph of the flow channel features. Scale bar, 5mm. **c)** Parametric sweep of incoming sweat pH (pH_in_ = 4.5 – 7.0) showing predicted pad pH as a function of *k*_*eq*_ in the (i) reaction pad and (ii) reference pad; solid curves denote the mean trend; dashed lines denote the predicted *k*_*eq*_ band. **d)** Colorimetric transduction and calibration. (i) Urease-catalysed urea hydrolysis increases local pH, shifting the phenol red colour and the RGB signal. (ii) Linear calibration of green-channel intensity (G) versus urea concentration (0–5 mM), with representative colour swatches. **e)** Smartphone readout workflow. (i) App interface with an on-screen overlay; blue and red circles register the reference and reaction pads, respectively, and dashed outlines indicate fiducial markers. (ii, iii) Representative readouts under cold and warm ambient lighting. **f)** On-body demonstration showing the patch worn on the forearm (i) and the corresponding app readout (ii). g) Comparison of quantitative urea measurements under no buffering, optimised microfluidic buffering and excessive buffering conditions using artificial sweat samples spiked with known urea concentrations.

By systematically varying p, ϕ and the incoming pH (4.5–7.0), we identified an optimal operating regime in which buffer release stabilises the reaction conditions (**Fig. 3c**). In the reaction pad (**Fig. 3c,i**), the final pH decreases with increasing buffer delivery rate (k_eq_), such that insufficient buffering results in elevated pH, whereas excessive buffering over-suppresses the enzymatic response. This behaviour defines a narrow window of k_eq_ that maintains the system within its effective operating range. In contrast, the reference pad (**Fig. 3c,ii**) remains stable near neutral pH across a broad range of k_eq_, providing a robust baseline for differential sensing. At the optimal design point (p = 0.45 mm, ϕ = 0.35), pH variation is minimised under all tested physiological conditions, stabilising the biochemical starting environment for urea sensing. These results demonstrate that controlled buffer release defines a physically constrained operating window, effectively decoupling sweat pH variability from sensor response.

To quantify urea through the buffered sensing chemistry, we first characterised the colourimetric response of the assay (**Fig. 3d**). Urease-catalysed urea hydrolysis increases local pH, inducing a colour shift of phenol red from yellow to red (**Fig. 3d,i**). This change was quantified using the green (G) channel of the RGB signal, which exhibited an approximately linear decrease with increasing urea concentration under controlled conditions (**Fig. 3d,ii**), enabling direct calibration for urea quantification.

To achieve reliable readout under practical imaging conditions, we implemented a smartphone-based analysis workflow (**Fig. 3e**). Fiducial markers were used to automatically locate the device and define a consistent region of interest, while image normalisation and colour correction were applied to reduce variations due to illumination and viewing geometry^24^. In addition, two identical sensing units were incorporated to provide internal redundancy and enable signal averaging. Using this integrated acquisition and correction strategy, the colourimetric signal could be robustly converted into sweat urea concentration^25,26^. On-skin measurements confirmed reliable device identification and stable colour capture across repeated acquisitions (**Fig. 3f**), demonstrating consistent readout under wearable conditions.

Having established the integrated microfluidic buffering and image-based quantification framework, we next evaluated the impact of buffering on quantitative accuracy by comparison to representative limiting conditions. To validate the necessity of controlled buffering, we compared the performance of our optimised microfluidic design with two limiting conditions using artificial sweat spiked with known urea concentrations (**Fig. 3g**). In the absence of buffering (“no buffer”, blue dots), the measured signal rapidly saturates, leading to significant overestimation and loss of quantitative capability. This condition mimics direct, unregulated sweat sensing (e.g., patch or strip-based measurements), where variations in sample composition distort the biochemical response. In contrast, under excessive buffering (purple dots), the signal becomes strongly suppressed and largely insensitive to urea concentration, as the reaction is over-neutralised. This reflects conventional bulk dilution approaches, where analytes are mixed into large volumes of buffer, compromising sensitivity. Only within the controlled buffering regime enabled by our structured microfluidic design does the measured signal exhibit a linear and accurate correspondence with the true urea concentration (red dots). This demonstrates that precise regulation of buffer delivery is essential to maintain the sensing system within its effective operating window.

### Data-driven translation of sweat urea into a clinically actionable blood urea

Having established a robust microfluidic buffering and colourimetric sensing framework for accurate quantification of sweat urea, we next asked whether these measurements could be translated into clinically meaningful estimates of systemic urea levels. A key challenge arises from the mismatch between sweat and blood urea concentrations. Although sweat urea may be proportionally related to blood urea, the precise relationship remains unclear, with reported ratios ranging from ∼1 to 50 depending on sampling protocols, physiological state and disease conditions^13,15,27^. This variability, driven by transport limitations and delayed equilibration between compartments, has hindered direct clinical interpretation of sweat measurements. Thus, following precise and standardised quantification enabled by our device, a reliable framework is required to infer blood urea from sweat-derived signals.

To address this, we performed a prospective clinical study and developed a data-driven calibration strategy that integrates sweat measurements with patient-specific information (**Fig. 4**). First, a training cohort from multicentre consisting of 129 patients and 475 paired measurements was used to construct the model. In the training phase, the calibration framework integrates heterogeneous data streams into a unified feature representation (**Fig. 4a**). Primary measurements (P), including sweat and blood urea, are combined with temporal information (T) and auxiliary clinical variables (A), including demographic parameters (age, sex and body mass index), renal-related indicators (renal replacement therapy status and chronic kidney disease history), and clinical severity indices (for example, APACHE II, SOFA and GCS scores). In addition, historic measurements are used to generate extended features (E) via linear prediction, enabling the model to capture temporal trends and delayed physiological responses. These components are concatenated into a feature matrix (X) for each patient (n_p_), where k denotes the number of samples per patient. Importance sampling and cross-validation are then applied to ensure robust model training.

**Fig. 4.**
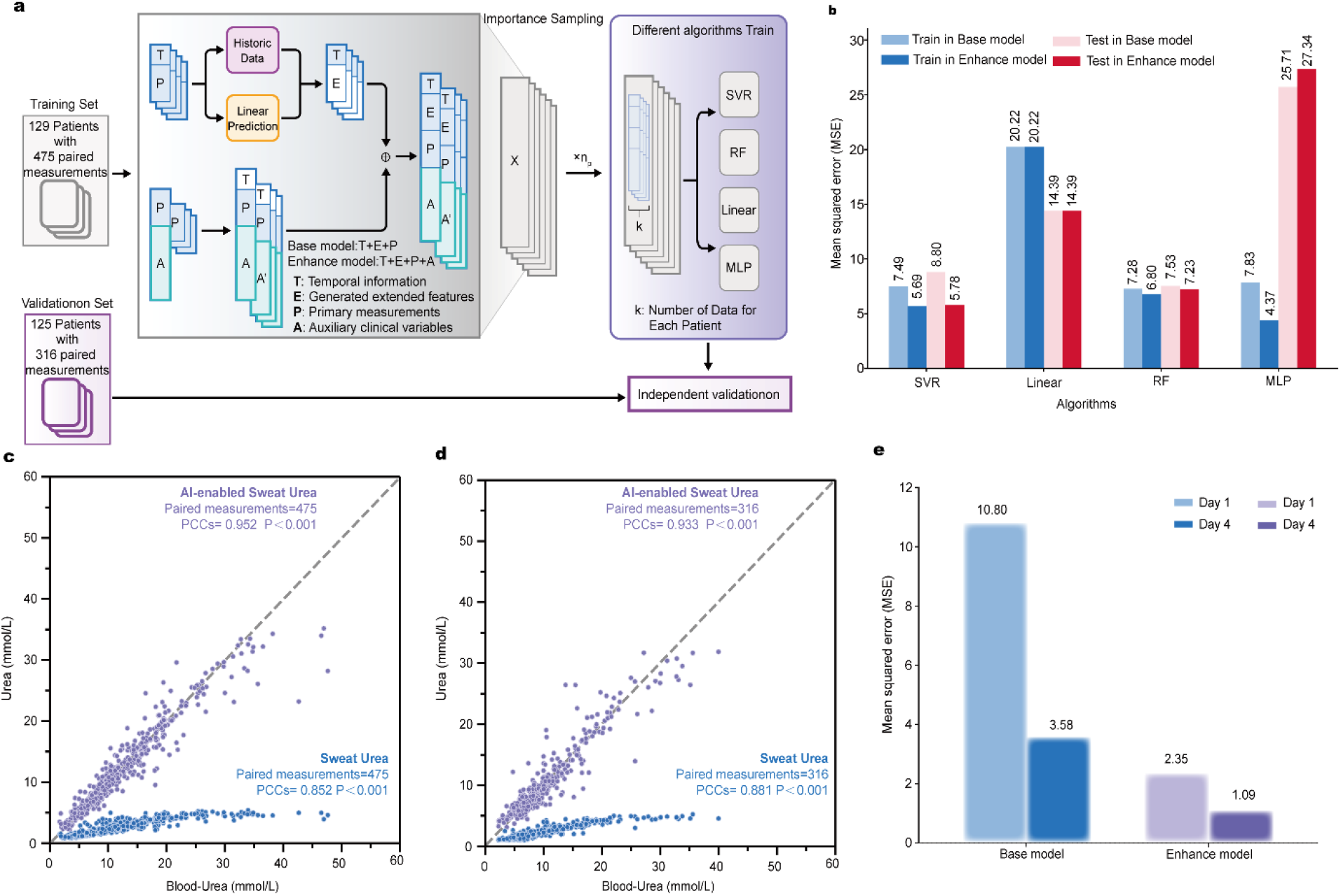
Development and validation of the AI-enabled calibration model for blood urea estimation. **a)**Schematic overview of the AI-estimated blood urea from sweat urea concentration model’s workflow. **b)** Mean squared error of different algorithms from two models. **c)** Correlation analysis between AI-estimated blood urea and blood urea across the training cohort. **d)** Correlation analysis between AI-estimated blood urea and blood urea across the validation cohort. **e)** Mean squared error on day 1 and on day 4 with the preceding day’s sweat and blood urea measurements incorporated as additional temporal inputs in different model.

Within this framework, two models were constructed to reflect distinct clinical scenarios. Base model is trained using only primary measurements (sweat and blood urea), corresponding to settings in which limited patient information is available and prediction must rely primarily on the sweat–blood relationship. In contrast, enhance model incorporates auxiliary clinical variables, enabling context-aware prediction when more comprehensive patient information is accessible. Notably, comparison with alternative algorithms, including linear regression, random forest, and multilayer perceptron, support vector regression (SVR) was selected as the calibration algorithm because it provides strong generalization for nonlinear relationships while maintaining a standardized and stable optimization framework (**Fig. 4b**). Consistent with this design, SVR-based enhance model achieved substantially lower prediction error than base model in the training cohort (mean squared error, 5.69 versus 7.49), demonstrating that integration of patient-specific physiological context improves model performance (**Fig. 4b**).

Using the trained SVR-based enhanced model, we then examined whether the framework could correct the nonlinear mismatch between sweat and blood urea. As shown in **Fig. 4c**, raw sweat urea measured from buffered microfluidics (blue) exhibits a positive but nonlinear relationship with blood urea (PCC = 0.852, P < 0.001), with progressive attenuation at higher concentrations. In contrast, AI-estimated blood urea (purple) shows a substantially improved agreement with ground-truth measurements (PCC = 0.952, P < 0.001), closely approaching the identity line across the full physiological range. The largest improvement occurs in the high-concentration regime, where raw sweat measurements deviate most strongly from linearity. These results demonstrate that the data-driven calibration effectively compensates for nonlinear distortion, physiological lag and interindividual variability.

To assess whether the trained model could generalise beyond the training dataset, we evaluated its performance on an independent validation cohort comprising 125 patients and 316 paired measurements. In this cohort, sweat samples were acquired using the same microfluidic device, while all patient data remained unseen during model training. This design enables a stringent test of whether the model can reliably translate sweat-derived signals into blood urea estimates in new patient populations and clinical environments. Using this independent dataset, predictions were generated from sweat measurements and compared directly with clinically measured blood urea values. Consistent with the training-phase observations, enhance model again outperformed the base model (mean squared error, 5.780 versus 8.800; **Fig. 4d**), yielding strong agreement with reference measurements (PCC = 0.933, P < 0.001), thereby confirming robust generalizability across populations and settings. Notably, incorporating prior-day sweat and blood measurements as additional temporal inputs further improved prediction accuracy (mean squared error, 10.80 versus 3.58 in the base model and 2.35 versus 1.09 in enhance model; **Fig. 4e**), approaching the accuracy of direct blood testing. This improvement highlights the benefit of integrating longitudinal information to better capture patient-specific physiological dynamics. These findings demonstrate that the combined microfluidic–AI framework enables accurate and generalizable estimation of blood urea from non-invasive sweat measurements.

### Renal risk stratification and clinical decision-making

Building upon the accurate estimation of blood urea, we next asked whether the AI-estimated blood urea from sweat urea concentration measured by buffered microfluidics could support clinically meaningful decision-making, in particular the identification and stratification of kidney injury. To increase statistical power for this analysis, we combined the training and validation cohorts into a unified dataset, leveraging all available paired measurements across centres.

To assess whether AI-estimated blood urea could support clinically relevant stratification, we examined how well it distinguished different stages of kidney injury across possible decision thresholds. The resulting receiver operating characteristic (ROC) curve yielded an area under the curve (AUC) of 0.953 for distinguishing AKI from non-AKI states (95% CI, 0.940–0.967; Fig. 5a), where an AUC of 1 indicates perfect separation and 0.5 indicates chance-level classification. Among patients with AKI, the AI-estimated blood urea also distinguished severe from non-severe disease with an AUC of 0.901 (95% CI, 0.873–0.929). Thus, despite the physiological differences between sweat and blood urea concentrations, the calibrated signal retained sufficient information to stratify both the presence and severity of AKI. To assess practical clinical performance, we examined classification outcomes at the optimal thresholds. For AKI prediction, the AI-estimated blood urea correctly identified 84.6% of measurements associated with AKI (true-positive rate) while accurately excluding 94.5% of non-AKI measurements (true-negative rate; **Fig. 5b**). For severe AKI, the model retained strong discriminative power, with corresponding true-positive and true-negative rates of 76.8% and 90.5%, respectively. This performance highlights the ability of the framework to both sensitively identify clinically significant kidney injury and reliably rule out low-risk cases using a non-invasive readout.

**Fig. 5.**
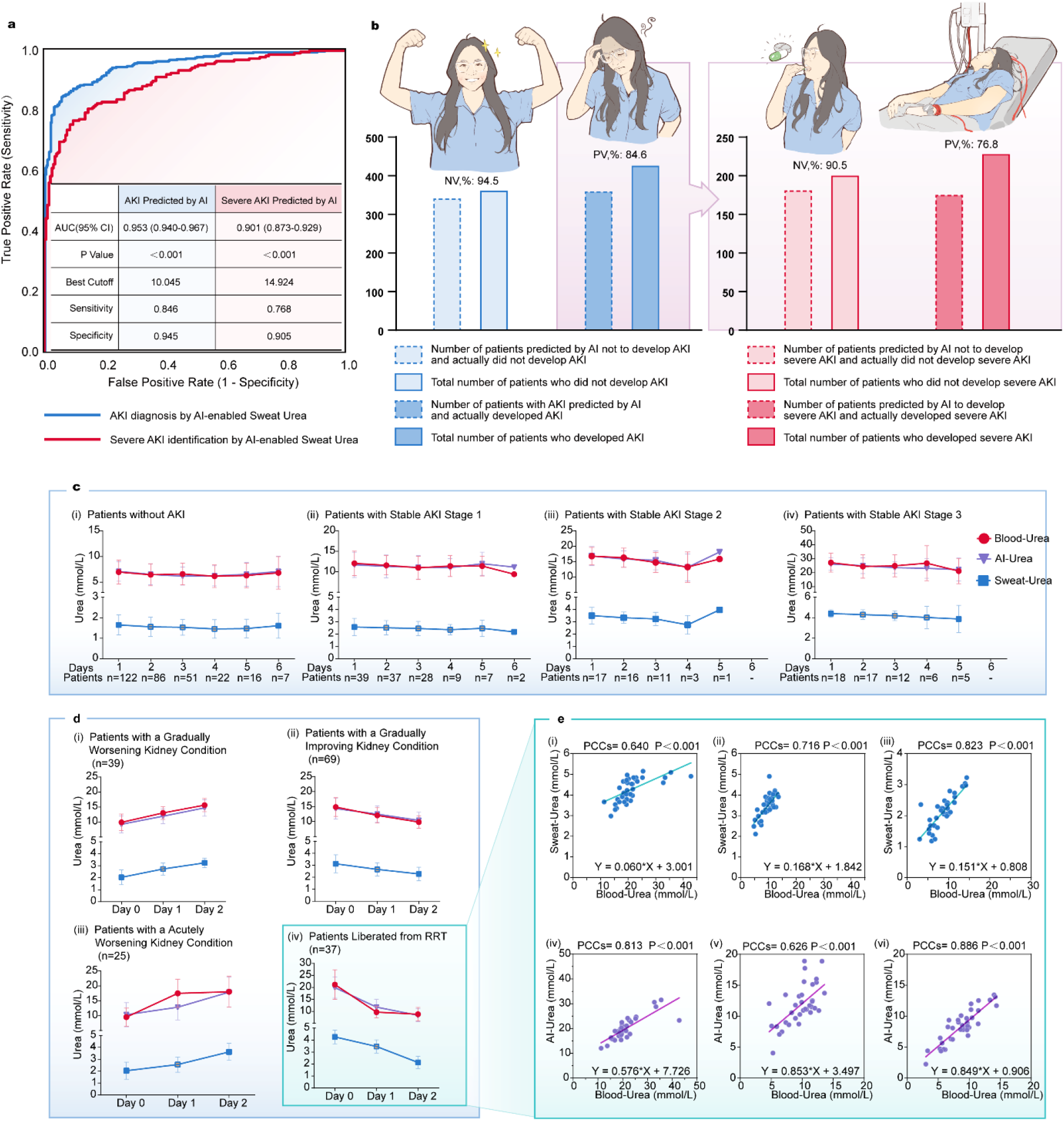
Clinical performance of the AI-estimated blood urea platform across kidney injury. **a)**Receiver operating characteristic (ROC) curves showing the ability of AI-estimated blood urea to distinguish patients with AKI and severe AKI, across different false-positive rates (1 − specificity) and true-positive rates (sensitivity). **b)** True-positive rate and true-negative rate of the AI-estimated blood urea model for AKI diagnosis (i), and severe AKI identification (ii). **c)**Dynamic performance of sweat urea, AI-estimated blood urea and blood urea in patients with stable kidney conditions stratified by AKI stage. ((i) Patients without AKI; (ii–iv) Patients with stage 1, stage 2, and stage 3). **d)** Dynamic performance of sweat urea, AI-estimated blood urea and blood urea in patients with changing kidney conditions. ((i) Patients with a Gradually Worsening Kidney Condition; (ii) Patients with a Gradually Improving Kidney Condition; (iii) Patients with an Acutely Worsening Kidney Condition; (iv) Patients Liberated from RRT). **e)** Daily dynamics for correlation and linear regression analyses between sweat urea and blood urea (i-iii), and between AI-estimated blood urea and blood urea (iv-vi), in Patients Liberated from RRT.

We next investigated whether the AI-estimated blood urea could capture physiologically meaningful variation beyond binary classification by examining longitudinal behaviour across patients with relatively stable clinical states. In individuals without AKI, as well as those with stable stage 1 and stage 2 AKI, sweat urea exhibited moderate correspondence with blood urea, while the AI-calibrated estimates closely tracked circulating levels with improved consistency over time (**Fig. 5c(i–iii)**). Notably, in patients with stable stage 3 AKI, buffered microfluidics measured sweat urea exhibited clear attenuation at higher concentrations, reflecting saturation of sweat urea at elevated systemic levels. In contrast, the AI-estimated values maintain monotonicity and better agreement with blood urea (**Fig. 5c(iv)**), indicating that the model compensates for nonlinear transport and exchange limitations in severe diseases.

We then assessed performance under dynamic clinical trajectories, where rapid physiological changes introduce temporal mismatch between sweat and blood compartments. In patients with gradually worsening or improving kidney function, both sweat urea and calibrated estimates showed consistent temporal trends that paralleled blood urea evolution (**Fig. 5d(i–ii)**). By contrast, under conditions of acute deterioration or following renal replacement therapy, blood urea changed abruptly, while sweat urea exhibited delayed responses, reflecting known equilibration lag between compartments. Importantly, the AI-estimated values closely followed the underlying blood urea trajectory despite these rapid transitions (**Fig. 5d(iii–iv)**), demonstrating that the model effectively mitigates temporal lag and captures dynamic physiological relationships.

To further examine how the calibration modifies the relationship between sweat and blood urea across conditions, we analysed the corresponding correlations before and after calibration (**Fig. 5e**). While sweat urea exhibited condition-dependent distortion and limited dynamic range, the AI-calibrated estimates showed substantially improved linearity and tighter agreement with blood urea across multiple clinical states. This behaviour indicates that the model not only improves numerical accuracy but also restores the underlying physiological mapping between compartments.

These results demonstrate that the AI-estimated blood urea not only enables accurate estimation but also provides clinically actionable information, enabling both accurate detection and stratification of kidney injury as well as continuous tracking of physiological dynamics. By integrating physicochemical stabilisation with data-driven calibration, this framework transforms sweat urea from a delayed and nonlinear surrogate into a real-time, non-invasive indicator of renal function, with potential utility for both screening and longitudinal monitoring in complex clinical settings.

## Conclusion

In this work, we establish an integrated microfluidic–AI framework for non-invasive assessment of renal function through sweat urea sensing and clinical translation. By combining a low-volume microfluidic device with controlled buffering and colorimetric readout, we achieve accurate and reproducible quantification of sweat urea using only microlitre-scale samples, while minimizing the impact of interindividual variability in sweat composition. The incorporation of structured fluidic buffering and optical standardisation enables stable assay conditions across heterogeneous physiological environments, providing high-fidelity measurements under real-world constraints. Building upon this measurement foundation, we develop a data-driven calibration strategy that translates sweat urea into clinically actionable estimates of blood urea by integrating patient-specific physiological context. This approach enables not only accurate estimation of systemic urea levels, but also robust renal risk stratification, including the detection and severity classification of AKI. Importantly, the framework captures nonlinear transport limitations and temporal lag between sweat and blood compartments, allowing physiologically informed tracking of renal dynamics across both stable and rapidly evolving conditions. Together, this work demonstrates that coupling physicochemical stabilisation with data-driven modelling can transform sweat-derived signals into reliable indicators of organ function, providing a scalable and patient-friendly alternative to conventional blood-based testing.

The integration of microfluidic sensing and artificial intelligence presented here highlights a broader opportunity for transforming accessible biofluids into clinically actionable information. Beyond urea, extension of this framework to additional biomarkers related to renal injury and systemic metabolism could further enhance diagnostic resolution and enable multidimensional assessment of disease states. The incorporation of longitudinal data streams and adaptive modelling strategies may also improve prediction accuracy across diverse patient populations and clinical scenarios, supporting personalised monitoring of disease progression and treatment response. More broadly, wearable systems that combine biochemical sensing with real-time data processing have the potential to evolve into continuous health monitoring platforms, capable of capturing dynamic physiological processes outside traditional clinical settings. By integrating biochemical measurements with complementary physiological signals, such platforms could provide a more comprehensive and context-aware understanding of health status. While the present study demonstrates near-clinical feasibility, its broader significance lies in establishing a generalisable strategy for translating complex peripheral signals into actionable clinical insights. This paradigm may ultimately enable earlier intervention, improved risk stratification and more responsive management of chronic and acute diseases.

## Methods

### 1. Microfluidic device design and moulding

A negative mould for the microfluidic channel layer was first fabricated by stereolithography-based 3D printing using a photosensitive resin (Formlabs Form 3, resin type V3, in-plane resolution 25μm). After printing, the mould surface was activated by oxygen plasma treatment (150 W power, 5 minutes duration) to promote subsequent silanisation. Immediately after plasma activation, the mould was vapour-phase silanised using 300 μL of a trichloro (1H, 1H, 2H, 2H-perfluorooctyl) silane reagent in a vacuum desiccator environment for 2 hours at 60°C temperature, thereby forming an anti-adhesive monolayer on the mould surface.

Polydimethylsiloxane (PDMS Sylgard 184) prepolymer and curing agent were mixed at a mass ratio of 15:1, degassed under vacuum (100mbar) for 3 mins, and cast onto the treated mould. The PDMS was then thermally cured at 60 °C for 6 h. After curing, the PDMS replica was carefully peeled off and rinse with IPA for 10 mins.

### 2. Hydrophilic surface modification of the PDMS microchannel layer

To render the demoulded PDMS microchannel layer hydrophilic, a grafting solution of methoxy-poly (ethylene glycol) silane (mPEG-silane, M_n_≈2 kDa) was prepared by dissolving mPEG-silane at 20 mg/mL in 95% ethanol. After neutralisation, a small amount of hydrochloric acid was added to adjust the solution to pH≈5. The solution was then introduced into the PDMS microchannels immediately after oxygen plasma activation (30W at 60s) by a syringe pump, filling the channel network and contacting all internal surfaces except the burst valve and outlet areas.

The hydrophilic treatment is attributed to (i) plasma-induced formation of surface silanol groups (Si–OH) on PDMS, which increases surface reactivity, and (ii) hydrolysis/condensation of the silane functionality, enabling covalent coupling (Si–O–Si linkages) of mPEG-silane onto the activated PDMS surface. The resulting PEG chains form a highly hydrated brush-like layer that reduces interfacial free energy and promotes water wetting, thereby producing a robust hydrophilic surface. After a dwelling time of 30 mins (static incubation, sealed), the channels were drained and the device was post-cured at 80 °C for 2 hours to promote silane condensation and stabilise the grafted layer. Following this process, the PDMS surface exhibited a water contact angle of 48° according to ASTM D7490-13.

Finally, the fabricated device was thoroughly rinsed with isopropyl alcohol (IPA) to remove residual reagents and was subsequently used for loading. Colorimetric porous film (0.45μm diameter,125μm from Shanghai Xinya) was employed to load the Phenol red as the pH indicator. The film was completely immersed in the phenol red solution (0.06 mg/mL in DI water) for 10 mins and subsequently baked to achieve complete drying under 50°C.

Buffer Tablets (pH 7 (Phosphate), Fisher Chemical™) conditioned to pH 6.8 to make up a volume of 100ml were used for pH measurement. 1mg urease reagent (grade 99%, solid, Sigma-Aldrich) was loaded into the elliptical reservoirs (long axis 1.6mm, short axis 1.1mm). For the encapsulation, a flat layer of PDMS without patterns was prepared using the same casting and curing procedure. After oxygen plasma treatment, the PDMS channel layer and the encapsulation layer were attached, and firm pressure was applied by parallel plates within 5 mins of surface activation to ensure complete bonding. Bonding was further stabilised by baking at 60°C for 1 h.

### 3. Depth-averaged phase-field model for microchannel filling

We consider a depth-averaged description of a shallow rectangular microchannel, where *h* denotes the unresolved channel height and ***u*** is the in-plane velocity averaged over that height. The scalar mask *H* is the phase-field indicator of the liquid phase, with *H* = 1 in the liquid and *H* = 0 outside the liquid. The mixture density and viscosity are denoted by *ρ*(*ϕ*) and *η*(*ϕ*), respectively. The shallow-channel drag term arises from the parabolic velocity profile across the channel height and is written as

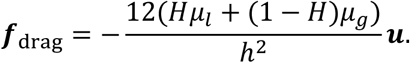

The unsteady depth-averaged Stokes equations are therefore

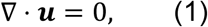

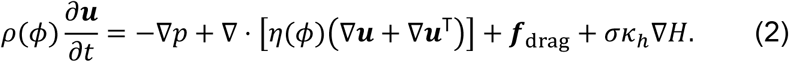

Here, *p* is the depth-averaged pressure and *σ* is the liquid–gas surface tension. The final term in Eq. (2) is the only wettability correction retained in the present model. It represents the out-of-plane capillary-pressure contribution induced by the contact angles at the unresolved top and bottom walls:

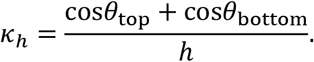

For identical top and bottom walls,

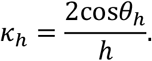

With the convention *H* = 1 in the liquid phase, the forcing *σκ*_*h*_∇*H* is localized near the diffuse liquid interface and accounts only for the thickness-direction contact-angle effect.

### 4. Coupled transport–reaction model for buffered urea sensing

A nonconservative Transport of Diluted Species (TDS) formulation was used to track the species across the phase field. This approach was adopted based on trade-off between computation costs and accuracy. The standard TDS equation for species *i* ∈ {*U, PBS, E, N, C*} is first written in conservative form as

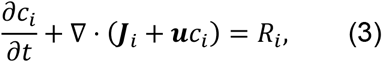

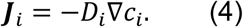

Here, *c*_*i*_ is the dissolved concentration of species *i*, ***u*** is the depth-averaged velocity, ***J***_*i*_ is the diffusive flux, *D*_*i*_ is the molecular diffusivity, and *R*_*i*_ is the volumetric production or consumption rate. The liquid mask is introduced to restrict diffusion, convection, and reaction to the liquid phase. The corresponding masked conservative form over the full computational domain is

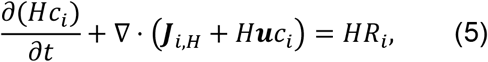

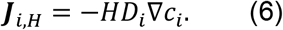

Thus, diffusion, convection, and reaction are active only where *H* ≃ 1. In the non-liquid region, where *H* ≃ 0, these transport and reaction terms are suppressed by the mask. Expanding the transient term gives

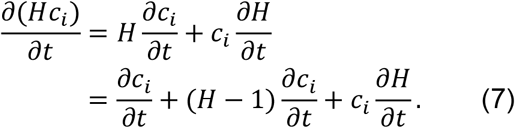

Substitution of Eq. (7) into Eq. (5) and moving the additional transient contributions to the right-hand side yields a TDS-compatible form in which ∂*c*_*i*_/ ∂*t* remains the primary time derivative:

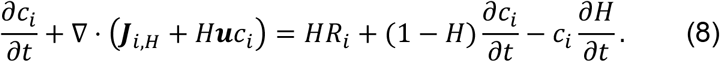

Equation (8) is algebraically equivalent to Eq. (5) when the two correction terms on the right-hand side are retained. These terms arise because the conservative masked equation evolves *Hc*_*i*_, whereas the TDS implementation solves directly for *c*_*i*_.

For incompressible flow Eq. (1),

the original TDS conservative equation, Eq. (3), is equivalent to the nonconservative form

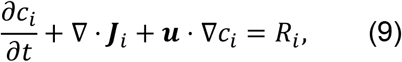

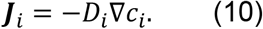

This equivalence follows from the expansion of the convective flux,

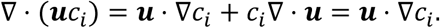

For the masked equation, the spatial variation of *H* introduces an additional contribution to the conservative convective flux:

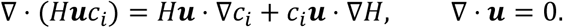

Therefore, the masked nonconservative form that remains equivalent to Eq. (5) is

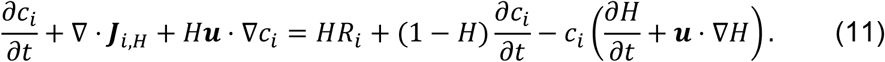

The last term contains the material variation of the phase-field mask,

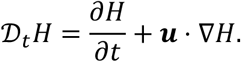

It is nonzero only where the diffuse interface evolves or is advected by the flow.

If the correction terms are neglected and the following simplified nonconservative equation is solved,

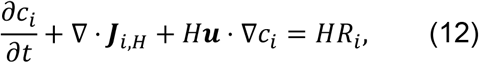

then the omitted source term is

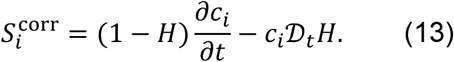

Equivalently, the residual with respect to the masked conservative equation is

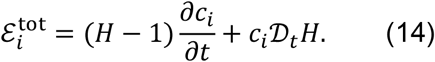

This residual can be decomposed into a transient masking error and a nonconservative advection error:

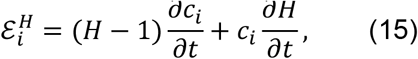

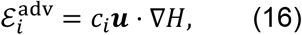

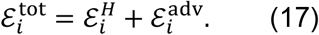

The first contribution results from replacing the conservative dependent variable *Hc*_*i*_ by *c*_*i*_. The second contribution results from using the nonconservative convective form in the presence of a spatially varying mask.

The corresponding global mass-balance residual is

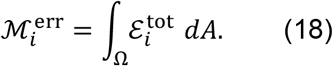

Let

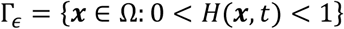

denote the diffuse-interface region of thickness *O*(*ϵ*). Since *H* = 1 in the liquid bulk and *H* = 0 outside the liquid phase, the residual is mainly localized inside Γ_*ϵ*_, provided that the concentration is fixed or inactive in the non-liquid phase. Hence,

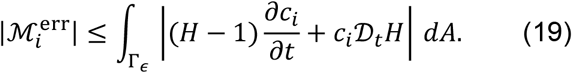

If the phase field is sufficiently narrow and the mask is transported consistently with the flow, so that *D*_*t*_*H* ≃ 0, then the integrated error decreases with the interfacial thickness:

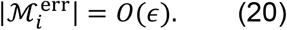

Therefore, the simplified masked nonconservative form is accurate when the diffuse interface is thin relative to the transport length scale and when the material variation of *H* remains small.

### 5. Surface dissolution of PBS

The PBS surface inventory *s*_PBS_(***x***, *t*) is defined on Ω_PBS_ and represents the amount of dissolvable PBS per unit surface area. Its depletion is controlled by the surface dissolution flux *J*_PBS_, the liquid mask *H*, a near-zero inventory gate ℋ_*δ*_, and a saturation gate *g*_sat,PBS_:

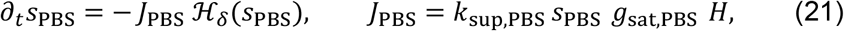

where *k*_sup,PBS_ is the surface supply rate in s^−1^. For a PBS recipe with molar fractions 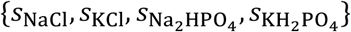, the limiting salt ∗ is selected as the least soluble component, and the saturation condition is evaluated using

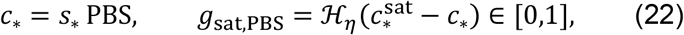

where 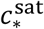 has units of mol m^−3^. The dissolved PBS released from the surface is converted into a volumetric source through the effective thickness *t*_*h*_ and the characteristic function 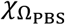:

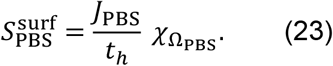

No separate PR concentration variable is introduced in this formulation.

### 6. Enzyme on the surface and in the bulk

On Ω_E_, the active surface enzyme density *s*_*E*_ (***x***, *t*) evolves through activation, desorption, and deactivation:

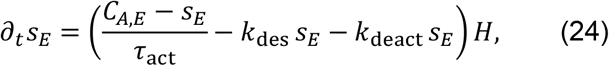

where *C*_*A,E*_ is the available surface enzyme loading, *τ*_act_ is the activation time, and *k*_des_ and *k*_deact_ are the desorption and surface deactivation rates. Surface Michaelis–Menten consumption and enzyme desorption are described by

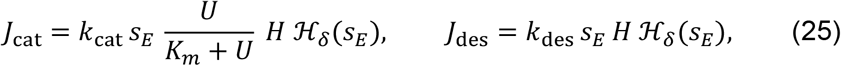

where *U* is the urea concentration, *K*_*m*_ is the Michaelis constant, and *k*_cat_ is the catalytic rate constant. These surface fluxes are mapped to volumetric sources in Ω_E_:

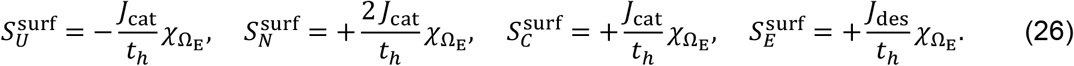

Here, *N, C*, and *E* denote the nitrogen-containing product, the carbon-containing product, and the dissolved enzyme concentration, respectively. Desorbed enzyme also catalyses urea in the liquid phase:

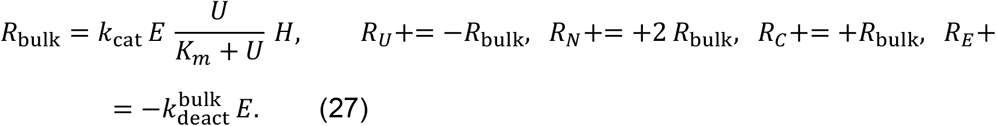

### 7. Recession of the solid and a release-rate correction

As PBS is consumed preferentially inside grooves, the solid–fluid interface can recede by a depth Δ, adding a quasi-stagnant diffusive film that acts in series with convection. A conservative correction is

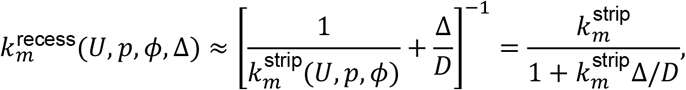

which captures the decay of cavity-floor transfer with increasing recess depth. In the calculations below, the effective groove recession depth is fixed at

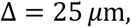

and the release coefficient used in the buffer-zone model is

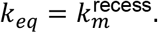

To optimize the reliability of the device, especially considering that the sweat of patients in clinical practice may fluctuate significantly due to the presence of lactic acid (pH range from 4.5 to 7.0), we calculated the effective surface mass transfer coefficient *k*_*eq*_ of the buffer zone for storing PBS based on additive resistance theory proposed in the literature,

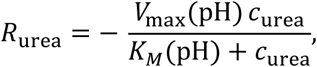

describing urease consumption of urea in the sensing pad, with *V*_max_ and *K*_*M*_ calibrated versus pH.

### 8. Image acquisition and urea concentration read-out

The correction pipeline assumes a linear radiometric domain such that multiplicative shading correction is physically meaningful. The application therefore acquires RAW sensor data (or an equivalent minimally processed stream that preserves linearity), including black-level subtraction and normalisation, and locks auto-exposure and auto-white-balance during capture.

Let Ω denote the ROI domain and let ***x***(***u***) ∈ *C* be the observed color vector at location ***u*** ∈ Ω, where *C* is the valid color space. Let *ℬ* denote the brightness reference region (enclosed by the device outline lines), {*Q*_*k*_}_*k*∈*K*_ the marker regions, and {*D*_*i*_}_*i*∈*J*_ the pad regions. The marker index set is *K* = {*c, α, β, γ, δ*}, where *Q*_*c*_ is the central marker and *Q*_*α*_, *Q*_*β*_, *Q*_*γ*_, and *Q*_*δ*_ are peripheral markers. The central marker provides a local reference at the template interior, while the peripheral markers provide reference observations distributed around the analytical region according to the external sampling design. The observed regional color in sampling region *S* ⊂ Ω is defined by:

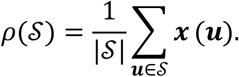

Brightness is represented by a scalar functional

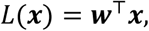

where ***w*** is a fixed channel-weight vector. The brightness reference value is computed from the brightness reference region as

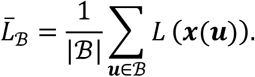

A global brightness gain is then estimated by comparing the measured brightness reference with a target brightness level:

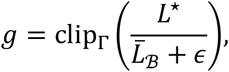

where *L*^⋆^ is the target brightness (defined as 170 in the clinical environment), *ϵ* is a numerical stabilizer, and Γ is the admissible gain interval. If temporal observations are available, the gain may be updated by a smoothed recurrence

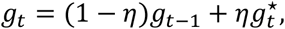

where 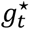 is the instantaneous gain and *η* is a smoothing coefficient. The brightness-normalized color field is

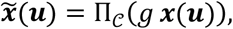

where Π_*C*_ projects the result into the valid color space. This first correction is global and is applied uniformly to all sampled regions.

After brightness normalization, each marker region is summarized as

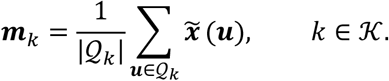

Each marker region has a known reference response ***a***_*k*_. Collecting the normalized marker observations and corresponding reference responses into matrices gives

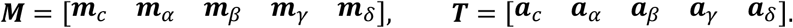

The color calibration matrix is estimated by regularized least squares:

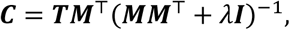

where *λ* is a regularization coefficient and ***I*** is the identity matrix. This matrix maps brightness-normalized observations to their reference responses in the least-squares sense. Each pad region is first averaged after brightness normalization,

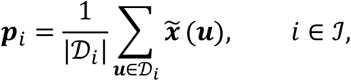

and is then calibrated by

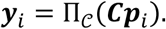

The green-channel intensity is then converted to urea concentration using the calibration function reported in the main text. For the dual-unit patch, the two inferred concentrations are compared for quality control; frames with disagreement greater than 10% are discarded, otherwise the averaged value is reported.

### 9. Machine learning via Support Vector Regression

Patient *i* is assigned to the test set with probability

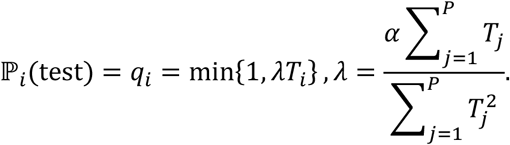

When truncation by min {1,·}is inactive for most patients; this choice yields

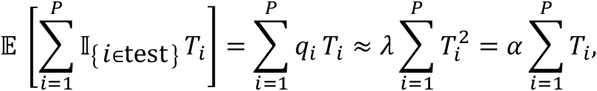

i.e., an expected record-level test fraction close to *α*, while ensuring *q*_*i*_ ≤ 1for long-stay patients. After splitting, each daily record contributes one supervised training pair (**x**_*i,t*_′ *y*_*i,t*_), where **x**_*i,t*_is the lag-embedded regressor defined above and *y*_*i,t*_is the measured blood-urea concentration on day *t*. For notational convenience in the optimisation, we re-index all training-set records by a single sample index *n* = 1, …, *N*, so that (**x**_*n*_^′^ *y*_*n*_)corresponds to some (**x**_*i,t*_′ *y*_*i,t*_).

Excluding a small subset of patient-identity–linked Primary Data (P Data) and explicit date/time descriptors (T Data), the remaining covariates are treated as conditionally exchangeable at the record level after lag embedding; the resulting feature dimensionality is modest. Nevertheless, incorporating T Data and additional Extra features (E Data) induces nonlinear interactions. We therefore adopt *ε*-SVR with a radial basis function (RBF) kernel:

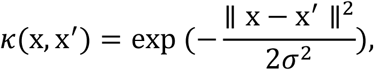

where *σ* > 0is the kernel length scale and ∥·∥denotes the Euclidean norm.

In the dual form, *ε*-SVR solves the convex quadratic program

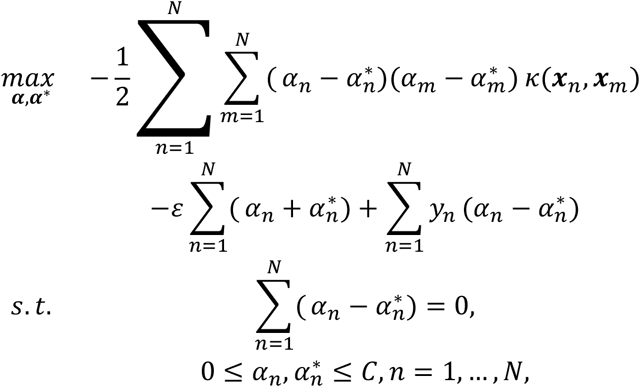

where *C* > 0is the penalty parameter controlling the trade-off between model flatness and violations of the *ε*-insensitive tube, and *ε* ≥ 0is the tube half-width. The resulting predictor is

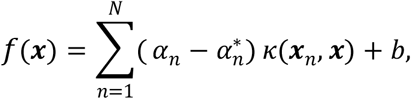

with *b* ∈ ℝthe bias term. Only samples for which 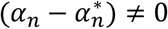act as support vectors and contribute to *f*.

### 10. Participants

This was a multicentre, prospective, observational study conducted at Zhongnan Hospital of Wuhan University, Shenzhen Second People’s Hospital, and Jingzhou Central Hospital. The study protocol was approved by the Ethics Committee (Approval No. 2025039K) in accordance with the principles of the Declaration of Helsinki. Written informed consent was obtained from all participants or their legally authorized representatives.

Inclusion criteria were as follows: aged >18 years; admission to the ICU for acute illness or post-operative care; and an anticipated ICU stay ≥24 hours.

Exclusion criteria included: (1) conditions preventing reliable sweat collection, including active skin lesions (e.g., wounds, burns, infection), severe scarring, or tattoos at forearm, or states of anhidrosis or marked hypohidrosis due to shock ; (2) key confounders potentially altering sweat composition, such as known systemic diseases affecting sweat secretion (e.g., cystic fibrosis, hyperhidrosis), chronic use (>4 weeks) of medications significantly impacting fluid/electrolyte balance (e.g., loop diuretics, thiazides, mineralocorticoid receptor antagonists), or the presence of significant pitting edema; (3) kidney transplant recipients; (4) a life expectancy of less than 48 hours, severe psychiatric or cognitive impairment precluding cooperation; (5) lack of informed consent.

### 11. Collection of paired sweat-blood measurements

Paired sweat-blood measurements were collected prospectively during routine ICU blood testing. For each predefined sampling event, sweat was collected from intact forearm skin using the wearable microfluidic patch immediately before a planned venous blood draw. After patch filling and colour development, the device was imaged using the smartphone readout system, and venous blood was drawn without delay. Blood urea was measured by the central laboratory at each participating hospital using routine automated clinical chemistry analysers. A paired record was defined as one valid sweat-urea measurement and the immediately subsequent blood-urea measurement from the same sampling event. Measurements with incomplete patch filling, failed image registration, visible leakage, image saturation or excessive disagreement between duplicate sensing units were excluded. Repeated paired measurements from the same patient were permitted during ICU admission. For each paired record, demographic characteristics, laboratory results, treatment information, renal replacement therapy status and kidney injury status were extracted from the hospital information system and linked to the corresponding sweat–blood measurement.

### 12. Statistical analysis

Continuous variables were reported as median with interquartile range or mean ± s.d., as appropriate. Categorical variables were reported as counts and percentages. Between-group comparisons were performed using the Mann–Whitney U test, Student’s t-test, χ^2^ test or Fisher’s exact test, as appropriate. Correlations were assessed using Pearson’s correlation coefficient. ROC curves were used to evaluate diagnostic performance, and AUCs with 95% confidence intervals were reported. A two-sided P value <0.05 was considered statistically significant.

## Declaration statements

### Data availability

The data that support the findings of this study are not publicly available due to restrictions imposed by institutional ethics committees and data governance policies of the participating hospitals. Access to the data may be considered upon reasonable request to the corresponding author and is subject to approval by the relevant ethics committees.

## Acknowledgements

P.Z. acknowledges the support from the National Natural Science Foundation of China (No. 82572484, No. 82272208). Y.Z. acknowledges the support from the Imperial’s start-up package and the UKRI Impact Acceleration Account. We also thank Shide Ren for the critical discussion.

## Author Contributions

J.Z., Z.S., X.R., Z.P. and Y.Z. designed the overall research framework. J.Z. and Z.S. contributed to device design, experimental implementation, data analysis and manuscript drafting. M.X., Y.G., G.L. and X.Z. contributed to microfluidic device fabrication, materials preparation, assay optimisation and device characterisation. S.L., C.Y. and M.L. contributed to clinical study coordination, patient recruitment and clinical data collection. G.P.M., Y.G. and P.M. contributed to experimental validation, data curation, statistical analysis and interpretation of results. Y.G., Z.S. and X.R. contributed to image-processing workflow development, model implementation and computational analysis. Z.P. supervised the clinical study design, clinical interpretation and multicentre coordination. Y.Z. and N.L. supervised the engineering design, analytical framework and overall project direction. J.Z., Z.S., X.R., Z.P. and Y.Z. wrote and revised the manuscript with input from all authors. All authors reviewed, edited and approved the final manuscript.

## Competing interests

The authors declare no competing interests.

## Notes

### Competing Interest Statement

The authors have declared no competing interest.

